# Improved yield of AAV2 and rAAV2-retro serotypes following sugar supplementation during the viral production phase

**DOI:** 10.1101/488585

**Authors:** Meghan Rego, Luke Makana Hanley, Ina Ersing, Karen Guerin, Meron Tasissa, Leila Haery, Isabelle Mueller, Erin Sanders, Melina Fan

## Abstract

Hyperosmolarity has been used as a means to improve yields in various processes such as antibody production in hybridoma cells and retroviral and adenoviral production systems. In this study we demonstrate that treatment of AAVpro 293T packaging cells with the sugars sucrose or sorbitol during adeno-associated virus (AAV) production improves the yields of AAV2 and rAAV2-retro serotypes 1.8-fold and 1.5-fold, respectively. Using an iodixanol gradient centrifugation strategy followed by a column-based buffer exchange, we show that the sugar supplements are not carried over to the final viral preparation and have no effect on the purity or infectivity of the virus. Given the wide availability, low cost, and ease of use of these sugars, we believe that this strategy can be easily adopted by vector cores and laboratories to improve yields of their AAV preparations.

## INTRODUCTION

In the early steps of viral infection, AAV2 particles attach to the cell surface via heparan sulfate proteoglycans before engagement and internalization by the protein receptor, AAVR (Summerford, 1998; Pillay, 2016). It is widely accepted that AAV2 yields are significantly lower than those of alternative serotypes and has been speculated that this is, at least in part, due to the affinity of AAV2 particles to surface heparin (Vandenberghe, 2010). During viral production, AAV2 particles remain tightly cellular associated limiting the amount of free virus in the supernatant and hindering the purification process. In an effort to improve yields of our AAV2 viral preparations we attempted to block the uptake of AAV2 viral particles by our packaging cells by inhibiting endocytosis.

Endocytosis is a critical cellular process by which external signals and nutrients are taken up by the cell. Often, cell surface proteoglycans serve as the initial attachment point for these molecules before being internalized by various endocytic pathways (Yanagishita, 1992; Hauser, 1999; Fuki, 2000; Fransson, 2004; Payne, 2007). Importantly, endocytosis can be inhibited by the addition of various compounds; for example, it has been demonstrated that hyperosmotic sucrose blocks clathrin-mediated endocytosis by sequestering clathrin microcages in the inner surface of the cell membrane thereby preventing clathrin adaptors from interacting (Heuser, 1989; Hansen, 1993). Interestingly, hyperosmolarity has also been used to improve protein production in various systems. When grown under hyperosmotic conditions, hybridoma cells display a reduced growth rate and enhanced monoclonal antibody production (Oyaas, 1989; Ozturk and Palsson, 1991; Oh, 1993). This phenomenon is not specific for antibody production as it has also been demonstrated during both adenoviral production in HEK293 cells and retroviral production in Te Fly A7 cells (Coroadinha, 2006; Shen, 2011). To our knowledge, this tactic has yet to be utilized to improve AAV production yields.

In this study, we examined the effects of sugar supplementation on the yields of AAV2, the AAV2 derivative rAAV2-retro and AAV5 viral preparations. We found that low levels of sucrose consistently improve yields without impacting cell viability. Interestingly, the effect of the sucrose supplementation varies across the serotypes tested with the largest improvement seen in the AAV2 and rAAV2-retro serotypes, 1.8-fold and 1.5-fold respectively, and no observable difference in our AAV5 batches. To assess whether this phenomenon was specific to sucrose or general to sugars we tested the effect of sorbitol on total yield and saw similar results. We believe that this supplementation should have no effect on downstream applications as sucrose or sorbitol were undetectable in the final purified viral preparations. As our study was limited to AAV2, AAV5 and rAAV2-retro it will be interesting to test this approach on the yield of other serotypes. Given the wide availability, low cost, and ease of use of these sugars we believe that this supplementation strategy can be easily incorporated by laboratories and vector cores to routinely produce AAV vectors with improved yields.

## MATERIALS AND METHODS

### Cell lines and Cell Culture Medium

AAVpro 293T cells were purchased from Clontech (Clontech 632273) and maintained in DMEM containing 4.5 g/L glucose, L-glutamine, and sodium pyruvate (Corning 10-013-CV) and supplemented with 10% heat-inactivated fetal bovine serum, FBS, (Seradigm 1500-500). Neuro2a cells were purchased from the American Type Culture Collection (ATCC CCL-13) and maintained in EMEM containing 1.5g/L sodium bicarbonate, non-essential amino acids, L-glutamine, and sodium pyruvate (Corning 10-009-CV) and supplemented with 10% heat-inactivated fetal bovine serum (Seradigm 1500-500).

### Plasmids

Plasmids for transfection were purified using Qiagen’s endotoxin-free HiSpeed Gigaprep kit (Qiagen 1054575) and quantified via spectrophotometry. The integrity of the cis plasmids and inverted terminal repeats (ITRs) were confirmed by restriction digest.

For the small scale viral preparations pAAV-CAG-GFP (Addgene 37825, gift of Dr. Ed Boyden) was used.

For the large-scale HYPERFlask M sucrose studies the following plasmids were packaged in the rAAV2-retro (Addgene 81070, gift of Dr. Alla Karpova) serotype; pGP-AAV-syn-jGCaMP7f-WPRE (Addgene 104488, gift of Dr. Douglas Kim) AAV-pgk-Cre (Addgene 24593, gift of Dr. Patrick Aebischer) pAAV-hsyn-Jaws-KGC-GFP-ER2 (Addgene 65014, gift of Dr. Ed Boyden) and pAAV.Syn.Flex.NES-jRGECO1a.WPRE.SV40 (Addgene 100853, gift of Dr. Douglas Kim). The following plasmids were packaged in the AAV5 serotype; pAAV-EF1a-double floxed-hChR2(H134R)-EYFP-WPRE-HGHpA (Addgene 20298, gift of Dr. Karl Deisseroth) pAAV-GFAP-hM3D(Gq)-mCherry (Addgene 50478, gift of Dr. Bryan Roth) pAAV-Ef1a-DIO eNpHR 3.0-EYFP (Addgene 26966, gift of Dr. Karl Deisseroth) and pAAV-hSyn-EGFP (Addgene 50465, gift of Dr. Bryan Roth). The following plasmids were packaged in the AAV2 serotype; pAAV-CAG-tdTomato (Addgene 59462, gift of Dr. Ed Boyden) and pAAV-hSyn-EGFP (Addgene 50465).

For the CellSTACK sucrose and sorbitol comparison studies pAAV-Ef1a-mCherry-IRES-Cre (Addgene 55632, gift of Dr. Karl Deisseroth) was packaged in the rAAV2-retro (Addgene 81070) serotype and pAAV-hSyn-DIO-hM4D(Gi)-mCherry (Addgene 44362, gift of Dr. Bryan Roth) was packaged in the AAV2 serotype

### Small scale viral preparation

6×10^6^ AAVpro 293T cells were seeded in T75 flasks and incubated overnight in DMEM supplemented with 10% FBS. The following morning, cells were transfected with a 1:1:1 molar ratio of transfer plasmid encoding GFP (pAAV-CAG-GFP, Addgene 37825) a plasmid encoding *rep* and the AAV2 *cap* genes and a plasmid encoding the adenoviral helper sequences. For transfection, 35 µg of total DNA per T75 flask was used. A 1:3 ratio by mass of total DNA to linear polyethylenimine (Alfa Aesar 43896-03) was prepared in serum-free OptiMEM (Thermo Fisher 31985-070) and incubated at room temperature for 15 minutes before adding to AAVpro 293T cells. Cells were maintained in DMEM supplemented with 2% FBS for the duration of the transfection, approximately 16-18 hours. The following morning, cells underwent a complete media exchange to DMEM supplemented with 4% FBS. Media was left untreated or supplemented with D-sucrose (MP Biomedicals IC821713). Cells were maintained in the sugar-containing media until harvest.

48 hours after transfection, cells and media were collected and cell viability assessed via trypan blue staining (Thermo Fisher T10282). Cells and media were sonicated on ice 3 times with a 50% amplitude and 1 second pulse using a Q125 Sonicator with 1/8” Diameter Probe (Qsonica Q125-110). The sample were then treated with 50 U/ml benzonase nuclease (Millipore 71205-3) for 45 minutes at 37°C. Following benzonase digestion, the samples were clarified via centrifugation at 3900 rpm for 10 minutes and the clarified supernatant collected and stored at 4° C.

### Large scale viral preparations

For sucrose studies viral vectors were prepared in HYPERFlask M vessels (Corning 10030). For each condition, 2 HYPERFlask M vessels were seeded with 1.8 × 10^8^ AAVpro 293T cells each in DMEM supplemented with 10% FBS and incubated overnight. The following morning, cells were transfected with a 1:1:1 molar ratio of a transfer plasmid, a plasmid encoding *rep* and the serotype-specific *cap* genes and a plasmid encoding the adenoviral helper sequences. For transfection 1 mg of total DNA per HYPERFlask M vessel was used. A 1:3 ratio by mass of total DNA to linear polyethylenimine (PEI) was prepared in serum-free OptiMEM and incubated at room temperature for 15 minutes before adding to AAVpro 293T cells. Cells were maintained in DMEM supplemented with 2% FBS for the duration of the transfection, approximately 16-18 hours. The following morning, cells underwent a complete media exchange to DMEM supplemented with 4% FBS. Media was left untreated or supplemented with 0.1M D-sucrose. Cells were maintained in the sugar-containing media until harvest.

For sucrose and sorbitol comparison studies virus was prepared in CellBIND-treated 2-chamber CellSTACKs (Corning 3310.) For each treatment, 1 CellBIND-treated 2-chamber CellSTACK was seeded with 1.2 × 10^8^ AAVpro 293T cells in DMEM supplemented with 10% FBS and incubated overnight. The following morning, cells were transfected with a 1:1:1 molar ratio of a transfer plasmid, a plasmid encoding *rep* and the serotype-specific *cap* genes and a plasmid encoding the adenoviral helper sequences.For transfection 0.8 mg of total DNA per CellSTACK vessel was used. A 1:3 ratio by mass of total DNA to linear polyethylenimine (Alfa Aesar 43896-03) was prepared in serum-free OptiMEM (Thermo Fisher 31985-070) and incubated at room temperature for 15 minutes before adding to AAVpro 293T cells. Cells were maintained in DMEM supplemented with 2% FBS for the duration of the transfection, approximately 16-18 hours. The following morning, cells underwent a complete media exchange to DMEM supplemented with 4% FBS. Media was left untreated or supplemented with 0.1M D-sucrose or 0.1M D-sorbitol (JT Baker, JTV045-7). Cells were maintained in the sugar-containing media until harvest.

96 hours after transfection, cells and media were collected and separated via centrifugation for 15 minutes at 3900 rpm. The clarified media was filtered through a 0.45 µm polyethersulfone membrane and viral particles precipitated with 8% w/v Polyethylene glycol for 4 hours at 4°C. The media was centrifuged for 15 minutes at 3900 rpm and the viral pellets resuspended in cell lysis buffer (50mM Tris, 150mM NaCl, 2mM MgCl2, pH 8.5). The virus was treated with 50 U/ml benzonase nuclease for 45 minutes at 37°C and then stored at 4°C. The cell pellet was resuspended in cell lysis buffer sonicated on ice 5 times with a 50% amplitude and 1 second pulse using a Q125 Sonicator with 1/8” Diameter Probe (Qsonica Q125-110). The samples were then treated with 50U/ml benzonase nuclease (Millipore 71205-3) for 45 minutes at 37°C. Following benzonase digestion, the samples were clarified via centrifugation at 3900 rpm for 10 minutes and then combined with the virus harvested from the media and stored at 4°C.

### Iodixanol gradients and buffer exchange

15% iodixanol (Sigma D1556-250) was prepared in PBS containing 1 M sodium chloride, 2.7 mM magnesium chloride and 2 mM potassium chloride. 25% iodixanol was prepared in PBS containing 2.7 mM magnesium chloride, 2 mM potassium chloride and mg/ml phenol red. 40% was prepared in PBS containing 2.7 mM magnesium chloride and 2 mM potassium chloride. Phenol red was added to 60% iodixanol to a final concentration of 0.01 mg/ml.

Quick-Seal Ultra-Clear Tubes (Beckman Coulter 34426) were used for the density gradients. Density gradients were prepared using sterile needles as follows; 10ml of 15% iodixanol underlayed with 8ml of 25% iodixanol underlayed with 8ml of 40% iodixanol underlayed with 3ml of 60% iodixanol. Approximately 8ml of virus was carefully overlayed onto the iodixanol gradient and the remaining space filled with PBS before sealing. Gradients were centrifuged in a T70i rotor for 350,000g for 90 minutes at 10°C. The tubes were punctured with an 18G needle at approximately 2 mm below the interface of the 60% and 40% gradients and the virus collected in 1ml fractions from the 40% layer. Clean fractions were pooled, buffer exchanged and concentrated using 100 KDa Amicon Ultra-15 Centrifugal Filter (Millipore UFC910008). Concentrated viral particles were resuspended in PBS containing 0.001% pluronic acid and 200 mM sodium chloride and stored at 4°C.

### Titering and qPCR

AAV preparations were titered using PowerUp SYBR green technology (Thermo Fisher A25741) and primers targeting the ITRs; forward ITR primer, 5’-GGAACCCCTAGTGATGGAGTT, reverse ITR primer, 5’-CGGCCTCAGTGAGCGA (Aurnhammer, 2012). AAV titer (GC/ml) was calculated based off of a plasmid standard, pAAV-CAG-tdTomato (Addgene 59462) of known concentration.

### Sucrose and sorbitol assay

Levels of sucrose in the purified viral preparation were quantified using the Glucose and Sucrose Assay Kit (Sigma Aldrich MAK013-1KT) according to the manufacturer’s instructions. Levels of sorbitol in the purified viral preparation were quantified using the D-Sorbitol Colorimetric Assay Kit (Sigma Aldrich MAK010-1KT) according to the manufacturer’s instructions.

### Endocytosis assays

AAVpro 293T cells were seeded in 6 cm^2^ dishes in DMEM supplemented with 10% FBS. The following day, media was exchanged with fresh DMEM supplemented with 4% FBS with or without 0.1M D-sucrose. After 72 hours media was removed and cells were incubated with 25 µg/ml pHrodo dextran (Thermo Fisher P35368) for 15 minutes at 37°C. Following incubation, the dye was removed, cells washed with PBS and imaged immediately.

AAVpro 293T cells were seeded in 6 cm^2^ dishes in DMEM supplemented with 10% FBS. The following day, cells were transduced for 1 hour with CAG-GFP rAAV2-retro in the presence or absence of 0.1M D-sucrose. The virus was removed, cells washed with PBS and media replaced with DMEM supplemented with 10% FBS. 72 hours later GFP expression was assessed by direct fluorescence.

### Silver staining

1×10^10^ genome copies were loaded onto NuPAGE 4-12% Bis-Tris Protein Gels (Thermo Fisher NP0335BOX) and proteins separated by polyacrylamide gel electrophoresis (PAGE.) Proteins were then stained using the SilverXpress Silver Staining Kit (Thermo Fisher LC6100) and the stoichiometric ratios and abundance of the viral capsid proteins VP1, VP2 and VP3 analyzed.

### Transduction

7,000 Neuro2a cells were transduced with 1µl of purified virus in one well of a 96-well plate. GFP expression was assessed 96h post transfection.

## KEY RESOURCES TABLE

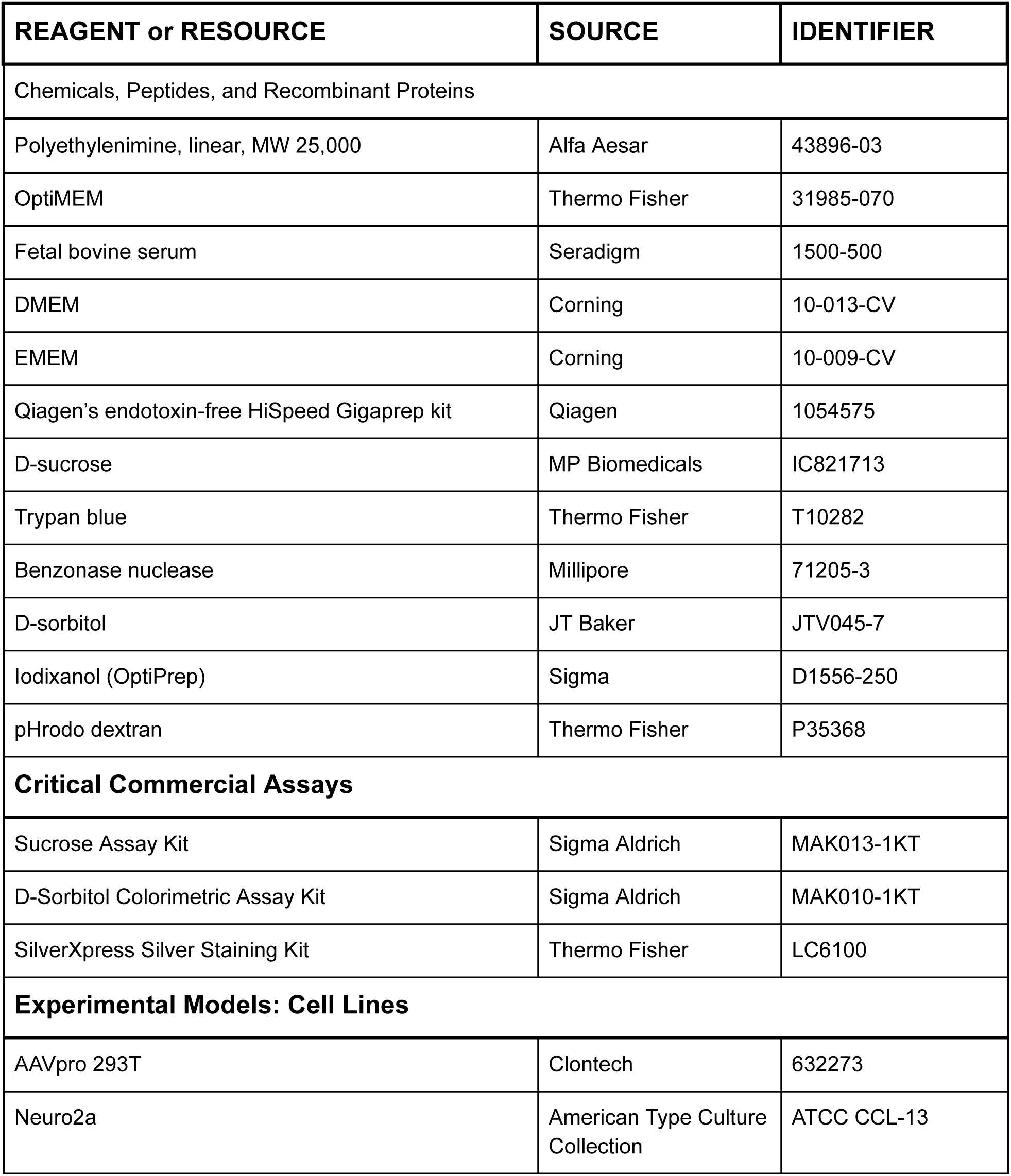
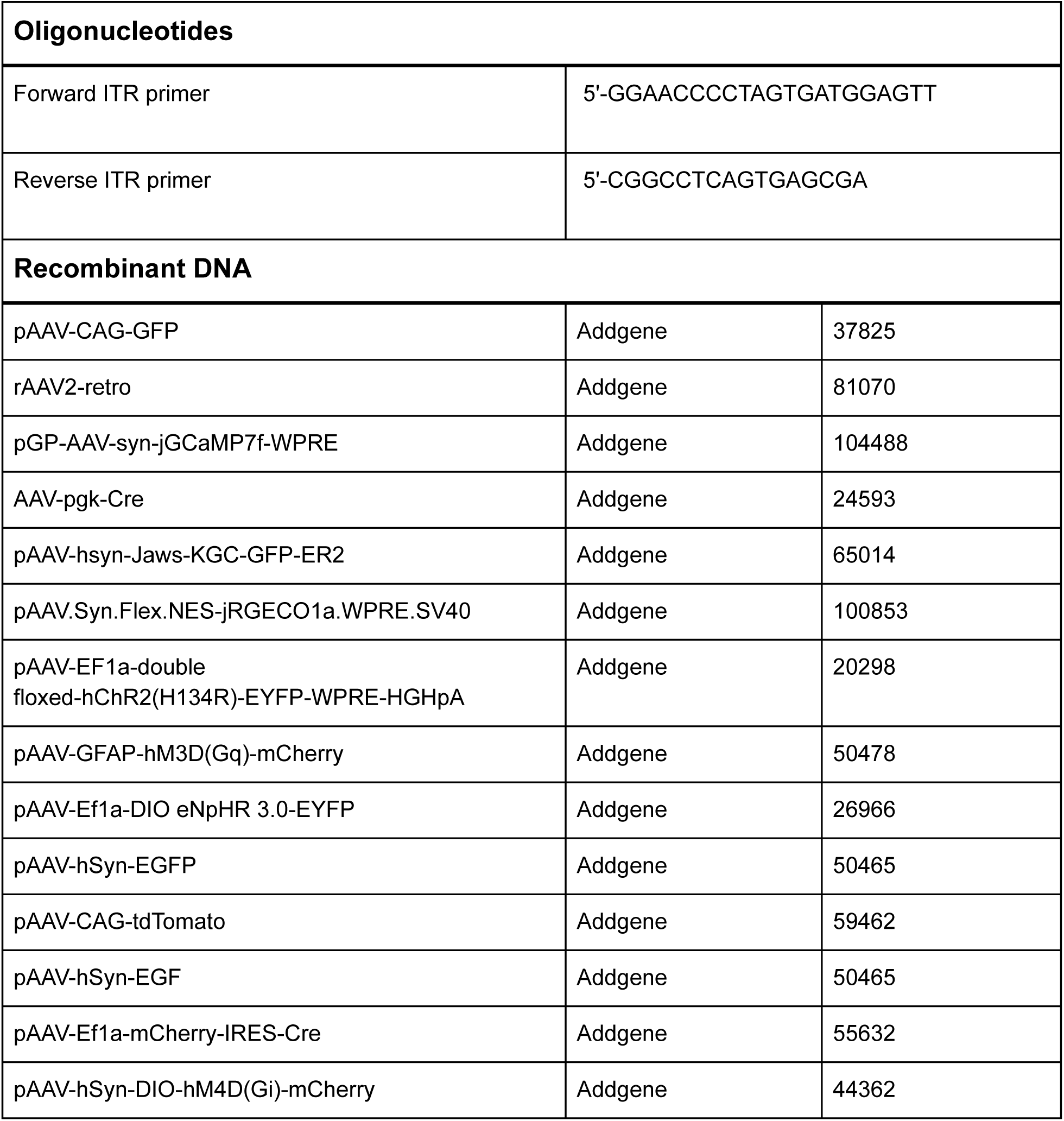

## RESULTS

### Low concentrations of sucrose improve AAV2 production without impairing cell viability

In an effort to improve our AAV2 titers we wanted to reduce the amount of viral uptake of our packaging cells via endocytosis. We reasoned that by blocking endocytosis, viral particles would remain in the cell culture supernatant thereby facilitating harvest and purification. While there are many chemicals available that inhibit endocytosis, we chose to pursue sucrose for several reasons; sucrose is relatively inexpensive, widely available, and unlike many of the other compounds that have been demonstrated to block endocytosis sucrose is not toxic and does not require special user handling.

To begin our studies AAVpro 293T cells were triple transfected with pAAV-CAG-GFP, a plasmid encoding *rep* and the AAV2 *cap* genes and a plasmid encoding the adenoviral helper sequences. We chose not to include sucrose in the media before or during transfection as PEI based transfection complexes use proteoglycans and various endocytic pathways for cell entry (Payne, 2007). The following morning, transfection mixes were removed and replenished with media only or media containing between 0 and 0.6M sucrose (Figure 1A and B) and incubated for 24 hours. Cells and media were then harvested and cell viability assessed via trypan blue staining. Viral vectors were isolated from both the cells and media, and the crude preparation was titered.

**Figure 1.**
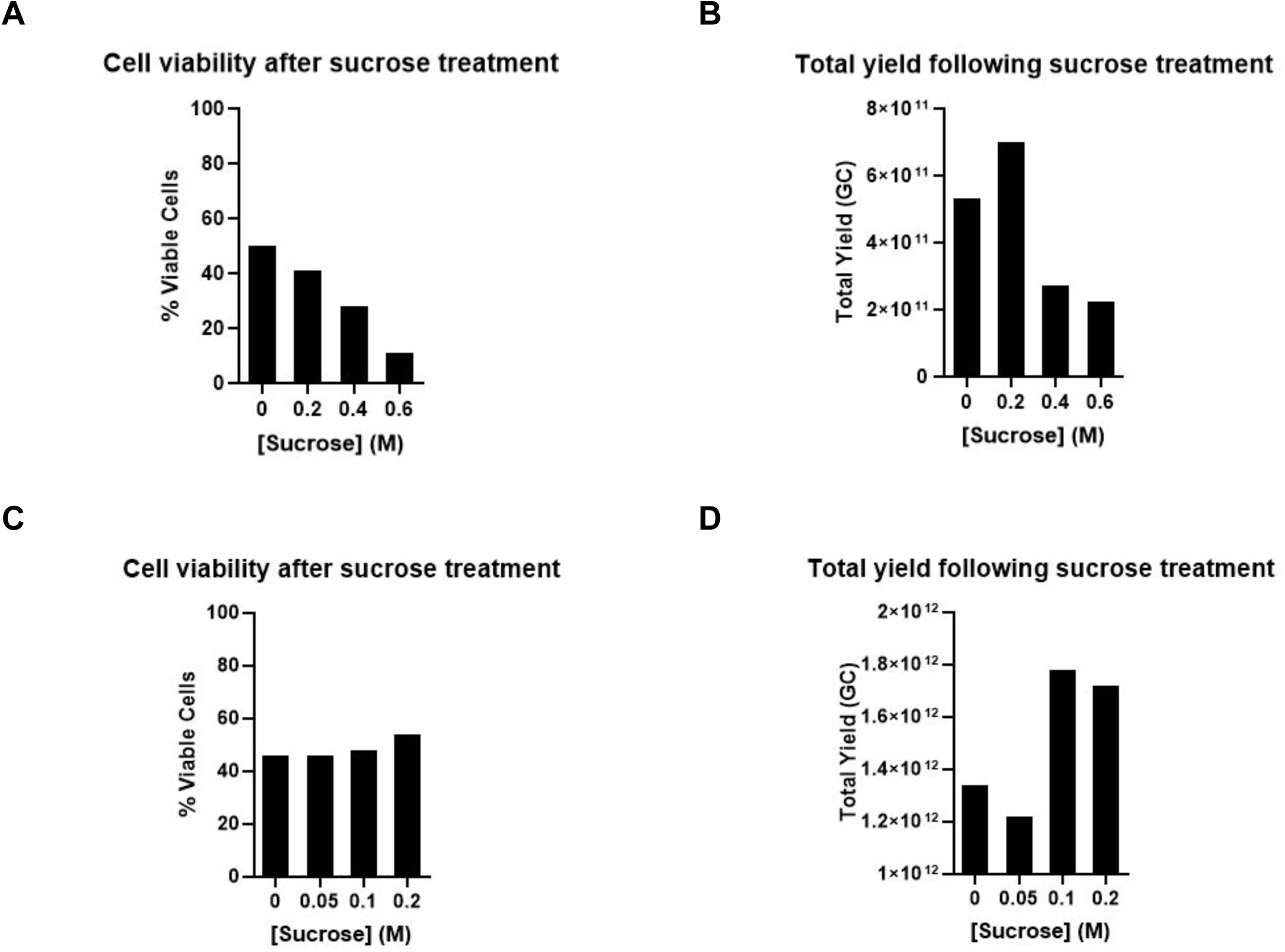
AAV yields after sucrose treatment in a small scale production. AAVpro 293T cells were triple transfected with a transfer plasmid encoding GFP, a plasmid encoding *rep* and the AAV2 *cap* genes and a plasmid encoding the adenoviral helper sequences. The following day media was replaced with fresh DMEM with or without sucrose. Cells viability was assessed 48h post transfection, A and C. Cells and media were then harvested and crude viral preparation titered, B and D.

The high doses of sucrose tested had a striking effect on cell viability (Figure 1A); at the time of harvest approximately 50% of the untreated cells were viable while only 28% and 11% survived after 0.4M and 0.6M sucrose treatment, respectively. At a dose of 0.2M, cell viability was slightly reduced to 41%. Interestingly, despite this drop in viable cells, we saw improved AAV2 yield in the 0.2M treatment group; indeed, there was a 1.3-fold increase in total AAV genome copies (GC) after low dose sucrose treatment (Figure 1B).

Given the cellular toxicity observed after treatment with high sucrose concentrations, we repeated the experiment with a range of low sucrose concentrations, 0-0.2M. At doses <0.2M cell viability was not affected (Figure 1C). When total yield was calculated, we saw an approximate 1.3-fold increase in GC in both the 0.1M and 0.2M sucrose treatment groups (Figure 1D). As increasing the sucrose concentration from 0.1M to 0.2M did not appear to improve yield, we chose to perform the remainder of our experiments at the lower dose.

### Variable effect of sucrose treatment on different AAV serotypes

We next sought to investigate how sucrose treatment would impact the total yields of our large scale purified viral preparations. For this study, 2 batches of AAV were prepared and purified and the total yield compared between untreated and sucrose treatment groups. AAVpro 293T cells were triple transfected with pAAV-CAG-tdTomato or pAAV-hSyn-EGFP, a plasmid encoding *rep* and AAV2 *cap* genes and a plasmid encoding the adenoviral helper sequences. The following day media was replaced with fresh DMEM with or without 0.1M D-sucrose and cells incubated for an additional 72 hours. Cells and media were harvested 96h post transfection and viral vectors purified via iodixanol gradient ultracentrifugation followed by concentration, buffer exchange and titering. Both batches had a substantially improved yield following sucrose treatment (Figure 2A). Indeed, we saw a 2.2-fold and 1.5-fold increase in total GC when preparing pAAV-CAG-tdTomato and pAAV-hSyn-EGFP, respectively (Figure 2B). Given these promising results, we decided to test the sucrose supplementation strategy on rAAV2-retro and AAV5 yields as these serotypes are routinely prepared and distributed by Addgene.

**Figure 2.**
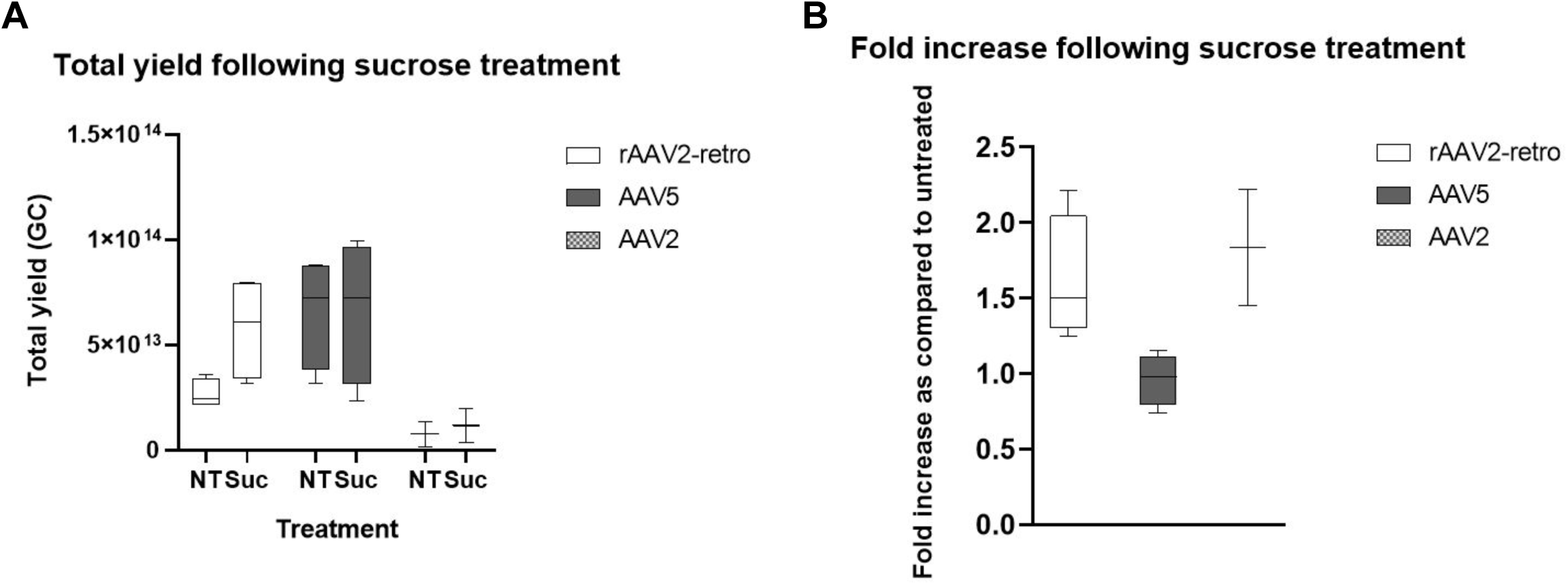
AAV yields after sucrose treatment in a large scale production. AAVpro 293T cells were triple transfected with a transfer plasmid, a plasmid encoding *rep* and serotype specific *cap* genes and a plasmid encoding the adenoviral helper sequences. The following day media was replaced with fresh DMEM with or without 0.1M sucrose. Cells and media were harvested 96h post transfection and viral particles purified via iodixanol gradient ultracentrifugation followed by concentration and buffer exchange. For each serotype between 2 (AAV2) and 4 (rAAV2-retro and AAV5) different preps were prepared and the range and average number of AAV genome copies (GC) calculated and plotted, A. B, the average fold change of sucrose to untreated viral prep was calculated and plotted.

For this study, 4 batches of AAV in the rAAV2-retro serotype and 4 batches of AAV in the AAV5 serotype were prepared and purified and the total yield compared between untreated and sucrose treatment groups. AAVpro 293T cells were triple transfected as described above. The following plasmids were packaged into the rAAV2-retro serotype; pGP-AAV-syn-jGCaMP7f-WPRE, AAV-pgk-Cre, pAAV-hsyn-Jaws-KGC-GFP-ER2 and pAAV.Syn.Flex.NES-jRGECO1a.WPRE.SV40. The following plasmids were packaged into the AAV5 serotype; pAAV-EF1a-double floxed-hChR2(H134R)-EYFP-WPRE-HGHpA, pAAV-GFAP-hM3D(Gq)-mCherry, pAAV-Ef1a-DIO eNpHR 3.0-EYFP, and pAAV-hSyn-EGFP. Not surprisingly, total yields varied significantly within the serotype-specific groups. The size of the expression cassette, tolerance of the packaging cells to the gene of interest, and natural variations between users and experiments will all influence the total yield. Despite the large spread of the data, however, there was a striking improvement in rAAV2-retro yields following sucrose treatment (Figure 2A). All 4 viral preparations saw between a 1.3 and 2.2-fold increase in yield with an average increase of 1.5-fold (Figure 2B). Intriguingly, we saw no difference between the untreated and sucrose-treated groups with the AAV5 serotype (Figure 2A and B).

### Improved yield is not specific to sucrose

We were curious whether the improved viral yields were specific to sucrose or caused by a general effect of the hyperosmotic conditions. To address this, we packaged pAAV-Ef1a-mCherry-IRES-Cre in rAAV2-retro and pAAV-hSyn-DIO-hM4D(Gi)-mCherry in AAV2. Following transfection, packaging cells were maintained in media only, media supplemented with 0.1M D-sucrose or media supplemented with 0.1M D-sorbitol. Cells and media were harvested 96h post transfection and processed as described earlier. We found that total AAV yields were improved in both serotypes regardless of the sugar used (Figure 3A and B).

**Figure 3.**
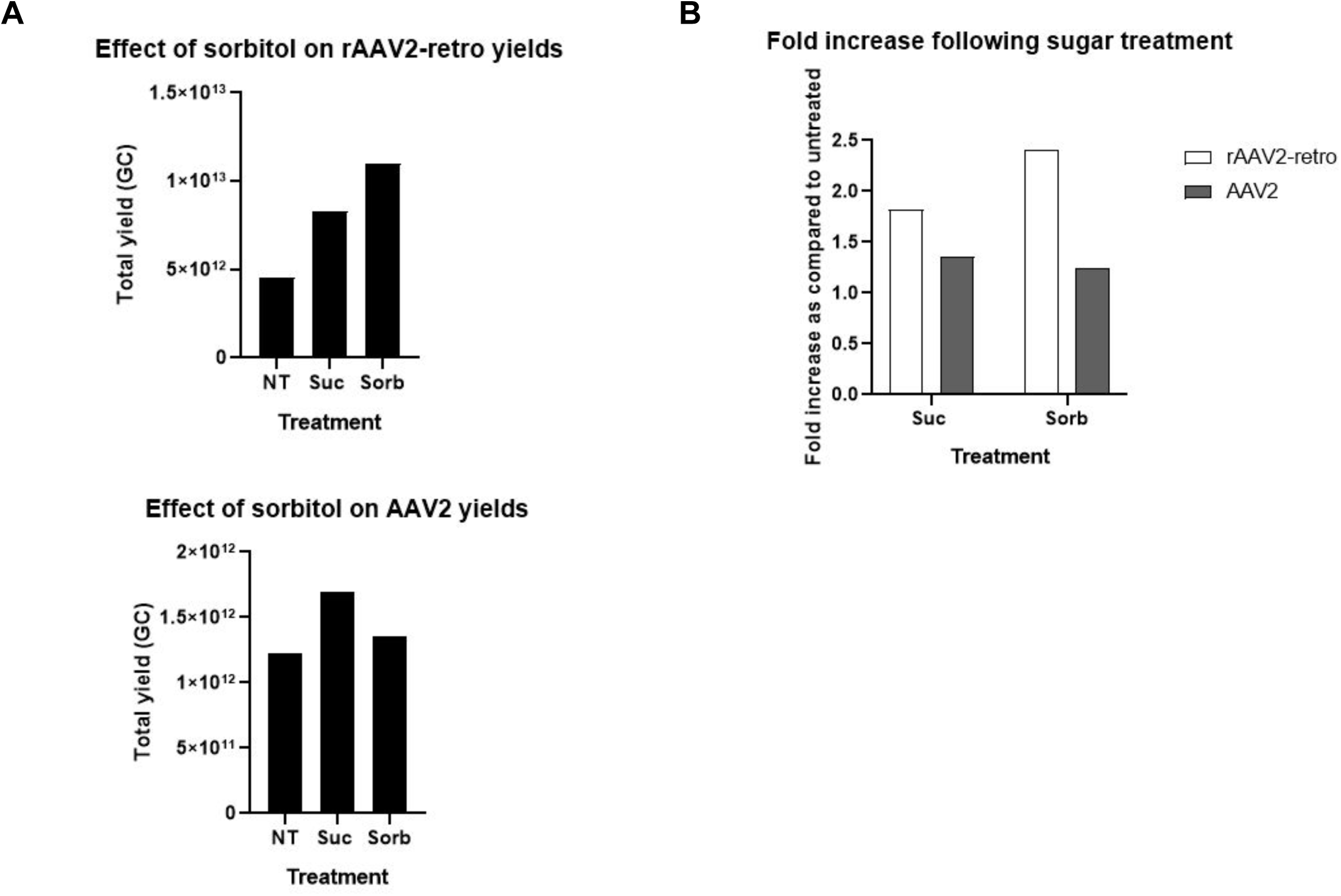
AAVpro 293T cells were triple transfected with a transfer plasmid, a plasmid encoding *rep* and serotype specific *cap* genes and a plasmid encoding the adenoviral helper sequences. The following day media was replaced with fresh DMEM with or without 0.1M of the indicated sugar. Cells and media were harvested 96h post transfection and viral particles purified via iodixanol gradient ultracentrifugation followed by concentration and buffer exchange. The number of AAV genome copies (GC) was calculated and plotted, A. B, the average fold change of sucrose to untreated viral prep was calculated and plotted.

### Sugar supplementation does not affect the purity or infectivity of the viral preparation

Addgene’s AAV preparations are used for a variety of applications including *in vivo* animal studies. We wanted to ensure that any changes we made to the viral production and purification process would not affect the final material. To assess this, we measured the sucrose and sorbitol concentrations in the final purified viral vector preparations as compared to the untreated preparations. Importantly, we were unable to detect any sucrose or sorbitol in the respective purified viral preparations (Figure 4A). Of note, 5% sorbitol is a commonly used excipient in AAV formulation buffers therefore trace amounts of the sugar are unlikely to be of concern.

**Figure 4.**
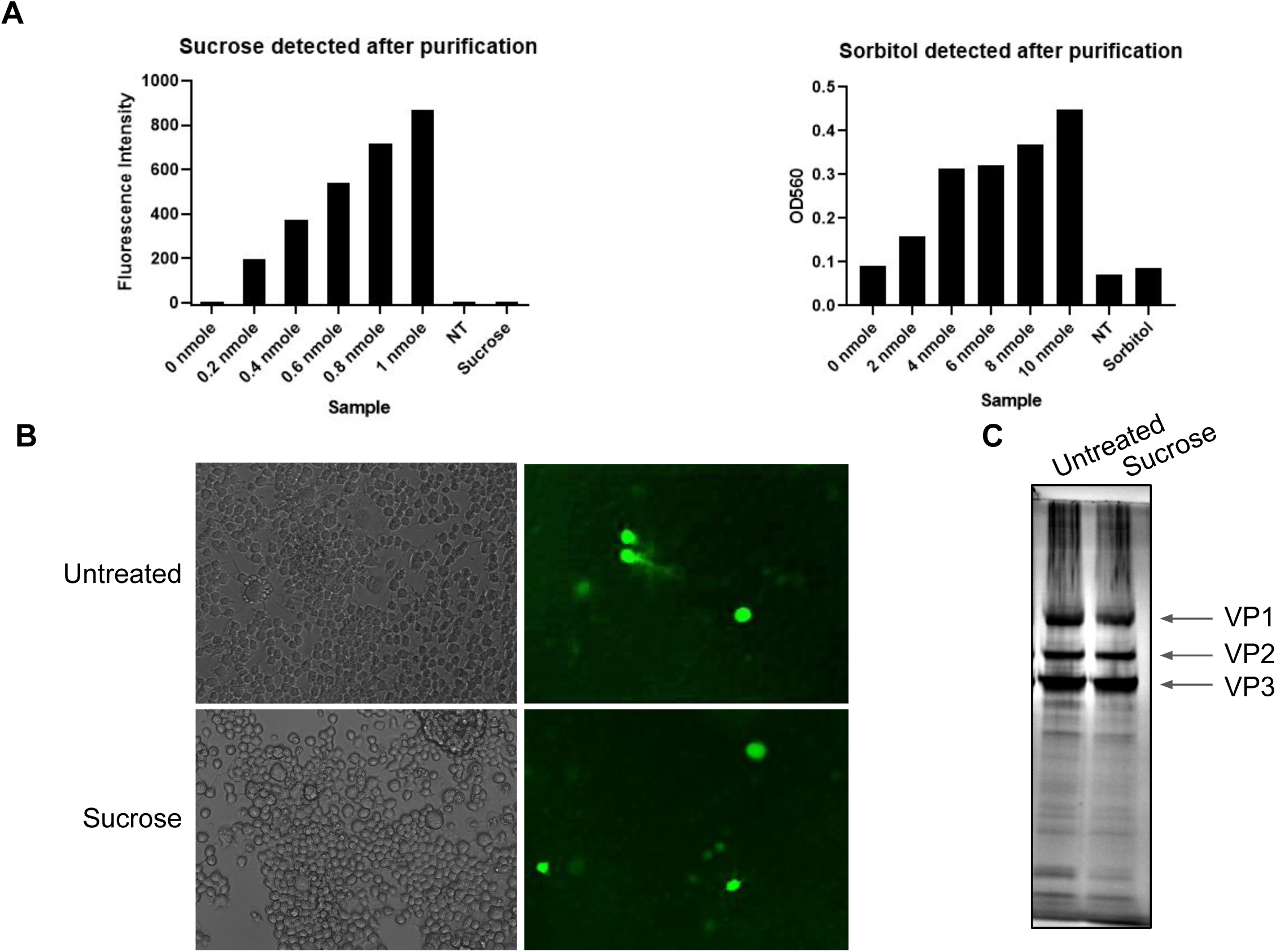
AAVpro 293T cells were triple transfected with a plasmid encoding jGCaMP7f (Addgene number 104488), a plasmid encoding *rep* and rAAV2-retro *cap* genes (Addgene number 81070) and a plasmid encoding the adenoviral helper sequences. The following day media was replaced with fresh DMEM with or without 0.1M of the indicated sugar. Cells and media were harvested 96h post transfection and viral particles purified via iodixanol gradient ultracentrifugation followed by concentration and buffer exchange. A, following buffer exchange, the level of sucrose or sorbitol was measured in the final preparations as compared to a standards of known concentration. B, Neuro2a cells were transduced with approximately 1.5 × 10^6^ vector particles per cell from the untreated and sucrose treated viral preparations. GCaMP7f expression was assessed 96h post transduction. C, purity of the final untreated and sucrose treated preparations was assessed via SDS-PAGE followed by silver staining.

To assess infectivity, Neuro2a cells were transduced with untreated or sucrose-treated pGP-AAV-syn-jGCaMP7f-WPRE rAAV2-retro and GCaMP7f expression analyzed 96 hours later (Figure 4B). Regardless of treatment, the GCaMP7f expression was easily observed indicating that the addition of sugars during packaging does not affect AAV function /ability to infect cells. To confirm that sugar treatment does not impact viral purity, viral proteins were separated by polyacrylamide gel electrophoresis (PAGE) and silver stained. The abundance of viral capsid VP1, VP2 and VP3 proteins compared to total protein levels was assessed as was the stoichiometric ratios of the capsid protein. We saw no difference in the purity of the viral preparation between treatments (Figure 4C).

### Prolonged sugar treatment does not impair viral transduction

We began testing sucrose supplementation with the assumption that sucrose treatment would block viral uptake by the packaging cells. To determine if this was indeed the case, we examined whether sucrose treatment blocks endocytosis of pHrodo green dextran dye. This dye is non-fluorescent outside of cells but emits green fluorescence when exposed to the acidic conditions of the endosome. AAVpro 293T cells were treated with 0.1M D-sucrose for 72h, stained with pHrodo green dextran dye and imaged immediately. Green fluorescence was observed in both treatment groups suggesting that endocytosis was functional (Figure 5A). In a parallel experiment, we tested the ability of AAV to infect in the presence of sucrose. AAVpro 293T cells were transduced with CAG-GFP rAAV2-retro in the presence or absence of sucrose and GFP expression assessed 72 hours later. Similar levels of GFP expression were observed between the treatment groups demonstrating that the presence of sucrose did not block viral uptake by the AAVpro 293T cells (Figure 5B). Interestingly, in a previous study investigating the mechanism of AAV entry by Stoneham et al. the group found that treatment with 0.4M sucrose did, in fact, block cellular transduction. These contradictory results could be due to the low concentration of sucrose used in our study or other experimental differences.

**Figure 5.**
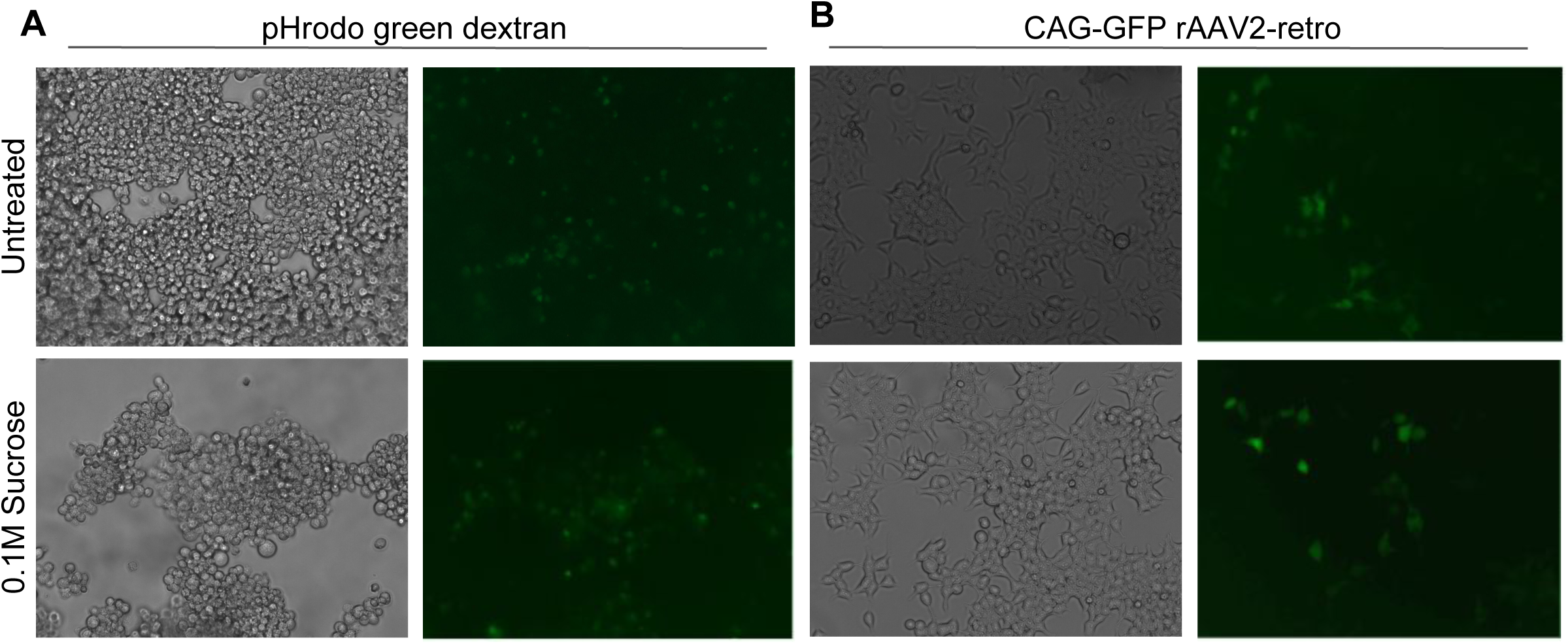
A, AAVpro 293T were left untreated (top panels) or treated with 0.1M sucrose (bottom panels) for 72 hours. The level of endocytosis of untreated and sucrose-treated cells was assessed via internalization of the pH sensitive pHrodo green dextran dye. B, AAVpro 293T cells were transduced with CAG-GFP rAAV2-retro in the absence (top panels) or presence (bottom panels) of 0.1M sucrose. 72h later, cells GFP expression was assessed by direct fluorescence.

## DISCUSSION

AAV production yields vary widely across serotypes owing to differences in the availability of viral particles in the cell culture medium versus those remaining in the packaging cells (Vandenberghe, 2010). The historically poor yields obtained from AAV2 serotyped viral vectors is thought to be due, at least in part, to the association and subsequent endocytosis of the viral particles to heparan sulfate proteoglycans on the surface of packaging cells. Previous work by Stoneham et al. demonstrated that treatment with 0.4M sucrose during AAV transduction significantly impaired viral uptake. We hypothesized that if AAV2 is indeed entering the packaging cells via HSPG-mediated endocytosis then we may be able to block this phenomenon by treating the cells with sucrose.

When producing AAV, we use a PEI-based transfection method; given that PEI utilizes endocytic pathways for cellular entry, we were concerned that pretreatment of cells with sucrose might impair our transfection efficiency. To prevent this, we chose to begin treatment after the initial transfection phase. In our initial small scale tests, we found that doses of sucrose >0.2M severely impaired cell viability and total viral yield. At dose <0.2M, however, we saw no effect on cellular viability and a modest but measurable improvement in AAV2 titers. We chose to pursue our studies with 0.1M D-sucrose as this dose resulted in the best overall yield without impacting cell viability.

The promising results of the small scale AAV2 study prompted us to examine how sucrose treatment would affect large scale AAV2, rAAV2-retro and AAV5 yields. Not surprisingly, on average we observed a 1.9-fold increase in our large scale AAV2 yields. Sucrose treatment also improved rAAV2-retro yields, albeit to a slightly lower level, 1.5-fold. Intriguingly, we did not see an effect of sucrose treatment on AAV5 yields. The different outcomes could be due to intrinsic properties of the different capsids. For example, rAAV2-retro may have similar results to AAV2 because, unlike AAV5, it was derived from AAV2 via directed evolution and may have retained many of the behaviors of the AAV2 serotype (Tervo, 2016). Alternatively, these differences may be an effect of overall yield. In general AAV5 viral preparations tend be high yielding while AAV2 and rAAV2-retro are much lower. Moreover, AAV5 has been reported to be released into the media during production, therefore blocking cellular uptake may not be critical for obtaining high yields (Vandenberghe, 2010). During the production of high yielding viruses such as AAV5, the packaging cells may be at or near protein production capacity. Consequently, the addition of sucrose has little to no effect on the final yield. In addition to sucrose, we demonstrate that sorbitol has a similar effect on viral yields. Importantly, neither sugar is carried over into the final preparation and sugar treatment does not impact the transducibility or purity of the final viral vector preparation.

To determine if these observations were indeed due to impaired endocytosis we examined the ability of our packaging cells to endocytose pHrodo green dye following sucrose treatment as well as their transducibility in the presence of sucrose. In both cases, we saw no difference in endocytosis between the untreated and sucrose treatment groups. While this seemingly contradicts the study by Stoneham et al. it should be noted that we are using about one fourth of the concentration of sucrose used in the previous study as this dose did not impair cell viability.

The osmotic stress response is a complex process involving the intersection of several different cellular pathways (Alfieri, 2007). While not completely understood several physiological changes could be contributing to the improved yield we are observing. It has been demonstrated, for instance, that in response to osmotic stress stationary phase cells have a 30-40% increase in RNA content resulting in a concomitant increase in protein production (Oh 1995, Lee 2000, Sun 2004). Moreover, under hyperosmotic conditions cells accumulate organic osmolytes that have been demonstrated to maintain cell volume, increase protein stability and refold misfolded proteins (Burg, 2007; Singh, 2011). The accumulation of these compatible osmolytes allows cells to adapt to mild osmotic stress and helps to build cellular tolerance to increased hypertonic stresses (Alfieri 2004).

While there is a strong precedent that the yield improvements we observed may be due to the osmotic stress response, we were not able to uncover direct evidence that this is the mechanism responsible. Of note, for our studies we only tested a single dose at a late time point. A thorough time course study examining the effects of different doses on a variety of readouts may be required to tease out the mechanism or mechanisms responsible.

Obtaining robust yields of AAV is a common goal among vector cores and laboratories routinely using AAV alike. We believe that media supplementation with sugars such as sucrose or sorbitol can provide these groups with an inexpensive and user-friendly method to improve yields of typically poor producing vectors. While we do not see an effect of sugar supplementation on our high yielding preparations, sugar supplementation has not proved detrimental; therefore, we believe that it could be incorporated into standard operating procedures as a fail-safe for unexpectedly poor preparations. Importantly, this study only examined the effect of sugar supplementation on AAV2, rAAV2-retro and AAV5 serotypes. It will be interesting to see if this strategy is effective when producing other serotypes as well.

## ACKNOWLEDGEMENTS

We thank Joanne Kamens and all other members of Addgene for advice, helpful discussion, and support during the preparation of this manuscript.

